# Population dynamics with threshold effects give rise to a diverse family of Allee effects

**DOI:** 10.1101/2020.04.02.021741

**Authors:** Nabil T. Fadai, Matthew J. Simpson

## Abstract

The Allee effect describes populations that deviate from logistic growth models and arises in applications including ecology and cell biology. A common justification for incorporating Allee effects into population models is that the population in question has altered growth mechanisms at some critical density, often referred to as a *threshold effect*. Despite the ubiquitous nature of threshold effects arising in various biological applications, the explicit link between local threshold effects and global Allee effects has not been considered. In this work, we examine a continuum population model that incorporates threshold effects in the local growth mechanisms. We show that this model gives rise to a diverse family of Allee effects and we provide a comprehensive analysis of which choices of local growth mechanisms give rise to specific Allee effects. Calibrating this model to a recent set of experimental data describing the growth of a population of cancer cells provides an interpretation of the threshold population density and growth mechanisms associated with the population.

## 1 Introduction

Mathematical models of population dynamics often include an Allee effect to account for dynamics that deviate from logistic growth (Allee and Bowen, 1932; Courchamp et al., 1999; Stephens et al., 1999; Taylor and Hastings, 2005). The logistic growth model can be written as

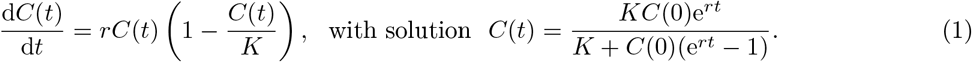

Here, *C*(*t*) ≥ 0 is the density at time *t*, d*C*(*t*)/d*t* is the growth rate, *r >* 0 is the low-density growth rate, *C*(0) is the initial density, and *K >* 0 is the carrying capacity density. Equation (1) has two *equilibria*: *C*^*^ = 0 and *C*\* = *K*, where an equilibrium is any value *C*\* such that d*C*(*t*)/d*t* = 0 when *C*(*t*) ≡ *C*\*. Since densities near *C*(*t*) ≡ *K* will approach *K*, while densities near *C*(*t*) ≡ 0 diverge away from zero, we say that *C*\* = *K* is a *stable equilibrium*, while *C*\* = 0 is an *unstable equilibrium*. This means that the logistic growth model implicitly assumes that all densities, no matter how small, eventually thrive, since 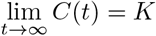 for *C*(0) > 0.

Mathematical models that include an Allee effect relax the assumption that all population densities will thrive and survive, which is inherent in (1) (Edelstein-Keshet, 2005; Murray, 2003; Stephens et al., 1999; Taylor and Hastings, 2005). Consequently, populations described using Allee effect models exhibit more complicated and nuanced dynamics, including reduced growth at low densities (Gerlee, 2013; Johnson et al., 2006; Neufeld et al., 2017) and extinction below a critical density threshold (Allee and Bowen, 1932; Courchamp et al., 1999; Taylor and Hastings, 2005). The phrase *Allee effect* can have many different interpretations in different parts of the literature. For instance, the *Weak Allee effect* is used to describe density growth rates that deviate from logistic growth, but do not include additional equilibria (Edelstein-Keshet, 2005; Murray, 2003; Stephens et al., 1999; Taylor and Hastings, 2005). A common mathematical description of the Weak Allee effect is

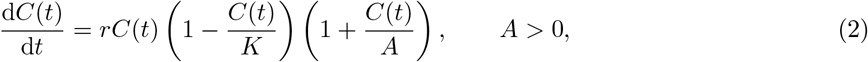

where the factor 1 + *C*(*t*)*/A* represents the deviation from the classical logistic growth model. Despite the similarity between (1) and (2), it is not possible to write down an explicit solution for (2) in terms of *C*(*t*), like we can for (1). Despite this, we are still able to examine the equilibria of (2) to understand the salient features of the Weak Allee effect. Since *A >* 0, (2) does not incorporate any additional equilibria other than *C*\* = 0 and *C*\* = *K*. Noting that the main feature of an Allee effect is a deviation from logistic growth, this cubic representation of the growth rate is employed predominantly for simplicity rather than explicit biological significance (Stefan et al., 2012; Stephens et al., 1999; Taylor and Hastings, 2005). Therefore, in this work, we refer to the Weak Allee effect as *any* population density growth rate that deviates from logistic growth without incorporating additional equilibria.

Another common type of Allee effect is the *Strong Allee effect*, in which an additional unstable intermediate equilibrium, *C*\* = *B,* with 0 < *B* < *K*, is incorporated (Courchamp et al., 1999; Edelstein-Keshet, 2005; Murray, 2003; Stephens et al., 1999; Taylor and Hastings, 2005). In such models, 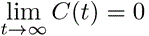 if *C*(0) < *B* and 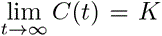 if *C*(0) > *B*, implying that *C*\* = 0 and *C*\* = *K* are stable equilibria and *C*\* = *B* is unstable. Typically, mathematical models incorporating a Strong Allee effect are written as

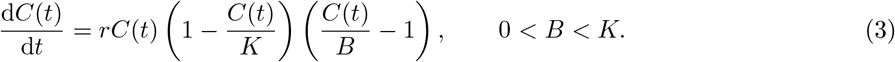

In a similar fashion to the Weak Allee effect, the cubic form of the Strong Allee effect is chosen predominantly for simplicity (Stefan et al., 2012; Stephens et al., 1999; Taylor and Hastings, 2005). Therefore, we will refer to any growth rate with two stable equilibria, *C*\* = 0 and *C*\* = *K*, and an additional intermediate unstable equilibrium as the Strong Allee effect. Throughout this work, we refer to growth rates that deviate from logistic growth as *an* Allee effect, whereas specific Allee effects (e.g., the Weak Allee effect and the Strong Allee effect) are referred to using more specific terminologies.

While Allee effects were originally used to describe population dynamics arising in ecology (Courchamp et al., 1999; Drake, 2004; Johnson et al., 2006; Seebens et al., 2017; Simberloff et al., 2013; Taylor and Hastings, 2005; Tu et al., 2019), there has been increasing interest in examining the potential for Allee effects in population dynamics relating to cell biology (Bobadilla et al., 2019; Böttger et al., 2015; de Pillis and Radunskaya, 2003; de Pillis et al., 2005; Gerlee, 2013; Jenner et al., 2019, 2018; Jin et al., 2017; Johnson et al., 2019; Johnston et al., 2017; Neufeld et al., 2017; Sarapata and de Pillis, 2014). In both cell biology and ecological applications, the Allee effect provides a suitable modelling framework to describe the dynamics of well-mixed populations that exhibit non-logistic features. However, because standard models incorporating Allee effects are continuum models that describe *global*, population-level features of the population dynamics, the interpretation of Allee effects at the individual scale remains less clear (Böttger et al., 2015; Johnston et al., 2017).

Understanding how local, stochastic growth mechanisms give rise to global Allee effects in a population is important, since these individual-level mechanisms can ultimately determine whether a population will survive or be driven to extinction (Böttger et al., 2015; Colon et al., 2015; Johnston et al., 2017; Scott et al., 2014). Certain individual-level biological features are ubiquitous among populations displaying Allee effects, providing a unifying feature in both cell biology and ecological applications. One of these phenomena is a *threshold effect* (Frankham, 1995; Metzger and Décamps, 1997; Rossignol et al., 1999), which we also refer to as a *binary switch*. We define a binary switch as a local feature of a population that behaves differently when a particular biological mechanism is present or absent. Some examples of binary switches include: the *go-or-grow* hypothesis in cell biology (Hatzikirou et al., 2012; Vittadello et al., 2020), phenotypic plasticity (Böttger et al., 2015; Friedl and Alexander, 2011), tree masting (Koenig and Knops, 2005), external harvesting pressure (Courchamp et al., 1999; Kuparinen et al., 2014), density-dependent clustering (Martínez-García et al., 2015), and resource depletion (Hopf and Hopf, 1985). For all of these examples, Allee effects have been proposed to potentially explain more complicated and nuanced population dynamics than are possible in a logistic growth framework. However, the link between the details of such a local binary switch and the resulting population-level Allee effect is unclear. Given that local binary switches are thought to be widely important in biology and ecology, we ask two questions: (i) how does the incorporation of a local binary switch in proliferation and death rates affect the global dynamics of a population? and, (ii) how does this local binary switch relate to different forms of Allee effects?

In this work, we show that incorporating local-level binary switches in a continuum, population-level mathematical modelling framework gives rise to a surprisingly diverse family of Allee effects. Some switches in proliferation and death rates give rise to established Allee effects, whereas other binary switches lead to more generalised Allee effects that have not been previously reported. We show that incorporating local-level binary switches in proliferation and death rates leads to a diverse family of Allee effects with only a few model parameters. This model, which we refer to as the *Binary Switch Model*, captures key biological features, but continues to exhibit the same qualitative features as various Allee effects. We conclude by applying the Binary Switch Model to a recent cell biology data set. Interpreting this data with our modelling framework suggests that the observed growth is non-logistic and that the phenomena is best explained by a binary switch at low density.

## 2 The Binary Switch Model

We consider an individual-based model framework that incorporates individual-level growth mechanisms varying with local population density to describe the temporal evolution of the global population density. One framework incorporating these aforementioned features is the stochastic agent-based model framework that we proposed in Fadai et al. (2019), in which individuals of the same size move, die, and proliferate on a two-dimensional hexagonal lattice. This discrete model incorporates exclusion (crowding) effects, allowing the population density to saturate at a finite capacity, as well as proliferation and death rates that vary with the local population density. While local population density can be measured in many different ways, Fadai et al. (2019) take the simplest approach and use the number of nearest neighbours as a measure of local density (Fig. 1).

**Figure 1:**
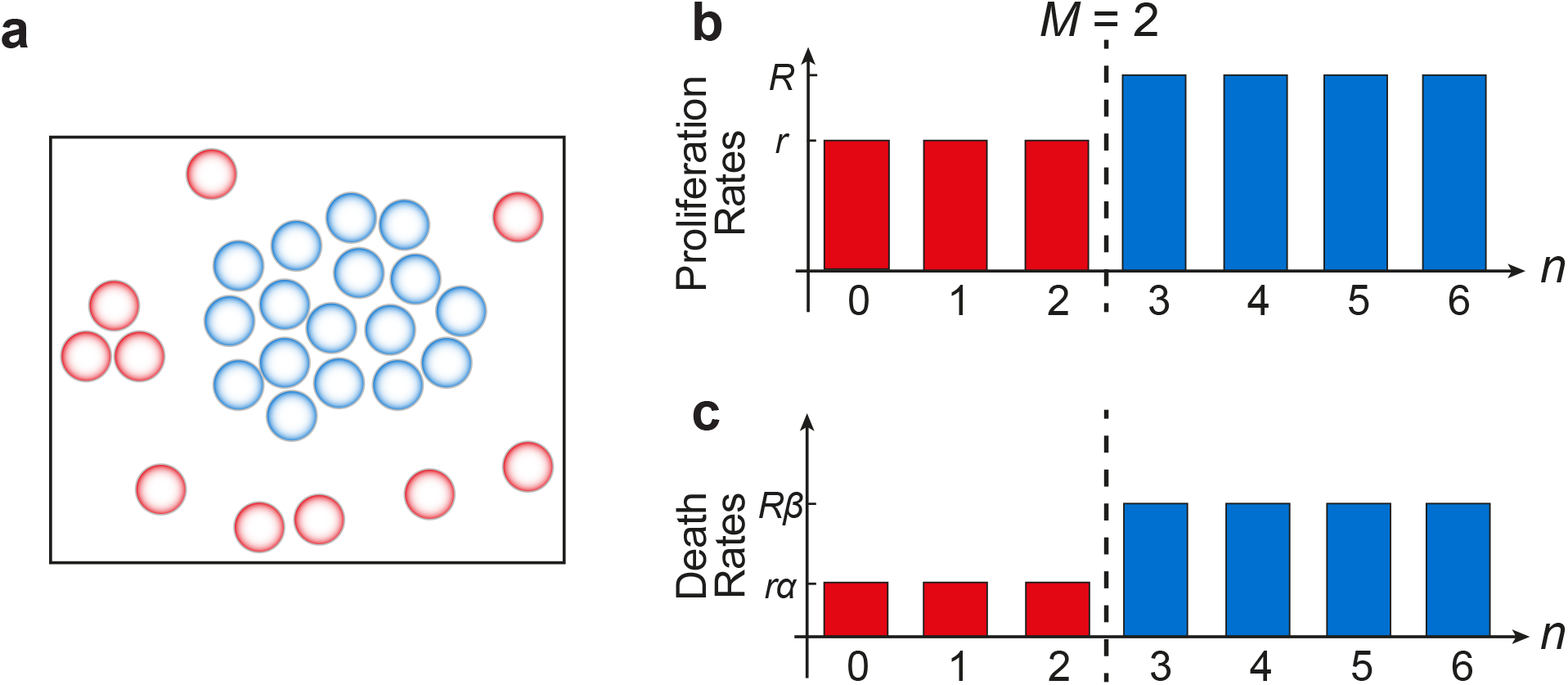
Schematic for the Binary Switch Model. Individuals in a population (a) can sense nearby individuals, providing a simple measure of local density. Individuals who sense higher than a threshold density, *M*, are shown in blue, while more isolated individuals are shown in red. This threshold density determines the constant rates at which individuals proliferate and die. (b,c) The binary switch shown here occurs when individuals can sense more than *M* = 2 neighbours.

As the individual dynamics of the stochastic agent-based model are difficult to analyse mathematically, we examine the continuum limit per-capita growth rate as a means of representing the average dynamics of the spatially uniform population, noting that there is good agreement between these two modelling approaches (Fig. 2). Full details of the discrete-continuum comparison is summarised in the Supplementary Information. Since the average population dynamics obtained from the discrete stochastic individual-based model agree well with its continuum description (Fig. 2), we will only consider the features of the continuum description of the model, whose per-capita growth rate is given by

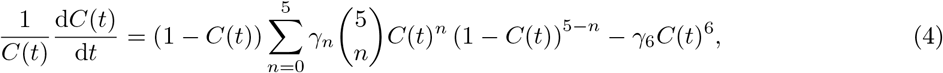

where

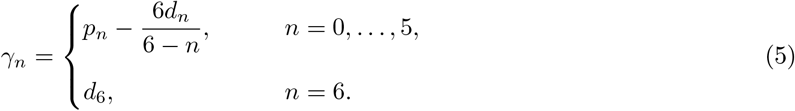

**Figure 2:**
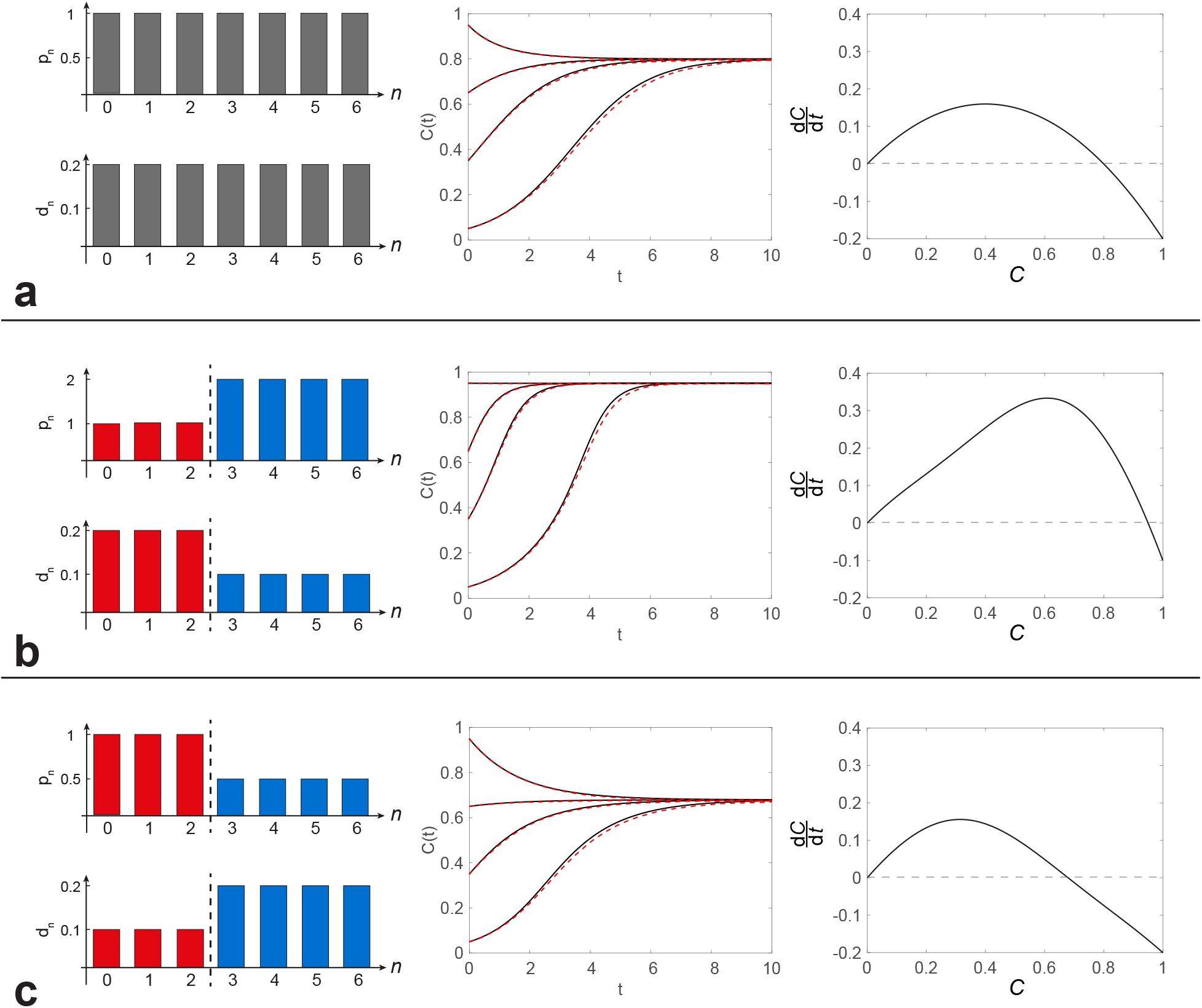
(a) When no binary switch is present, (4) reduces to logistic growth. (b,c) When a binary switch is incorporated in proliferation and death rates (*M* = 2), the continuum limit is no longer logistic. In all of these parameter regimes, the average density data determined from discrete model simulations, shown in red dashed curves in the middle column (Supplementary Information), agrees well with the continuum limit predictions (7), shown in black solid curves. Density growth rates in the right-most column show that (a) is logistic, while (b,c) are not.

Here, *C*(*t*) is the population density at time *t*, while *p_n_* and *d_n_* are the proliferation and death rates that vary with the number of nearest neighbours, *n* (Fadai et al., 2019). The parameter grouping *γ_n_* can be interpreted as the net growth mechanism for a particular local population density. Noting that *C*(*t*) ≡ 1 represents the maximum packing density, we have *C*(*t*) ∈ [0, 1]. Equation (4) has a thirteen-dimensional parameter space: namely, **Θ** = (*p*_0_*, …, p*_5_*, d*_0_*, …, d*_6_).

We incorporate a binary switch into (4) by choosing

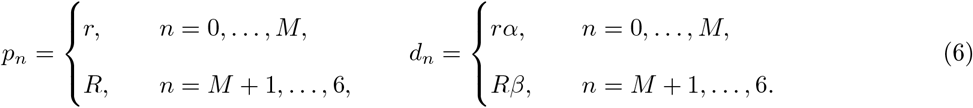

This choice of parameters means that we have the proliferation rate *p_n_* = *r* when the local density is at or below the critical density *M*, or *p_n_* = *R* when the local density is above *M*. We refer to *M* ∈ {0, 1, 2, 3, 4, 5} as the *threshold density*. For simplicity, we assume that the death rates are a particular fraction of the proliferation rates: *α* ∈ [0, 1] and *β* ∈ [0, 1]. It is useful to note that (4)–(6) relaxes to the classical logistic growth model, for any choice of *M* ∈ {0, 1, 2, 3, 4, 5} by setting *r* = *R* and *α* = *β* (Fig. 2a).

By substituting (6) into (4), we obtain the *Binary Switch Model*,

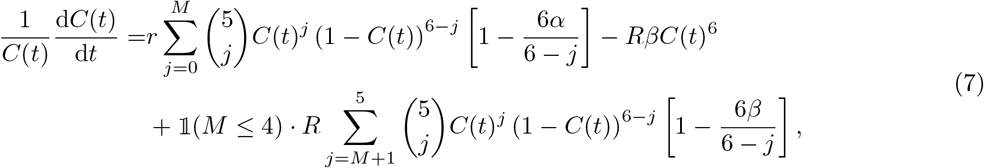

where

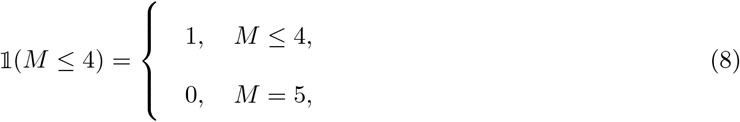

is an indicator function. The Binary Switch Model shows, for the first time, how a local binary switch in individual-level proliferation and death rates leads to a particular global density growth rate. A summary of parameters and their particular biological interpretation is shown in Table 1. In particular, we note that the Binary Switch Model reduces the thirteen-dimensional parameter space in (4) to a five-dimensional parameter space: **Θ** = (*r, R, α, β, M*). This reduced parameter space means that the Binary Switch Model can be used with less risk of over-fitting than (4) (Warne et al., 2019). We will discuss further merits of this reduced parameter space when calibrating the Binary Switch Model to experimental data in Section 3.

**Table 1:**
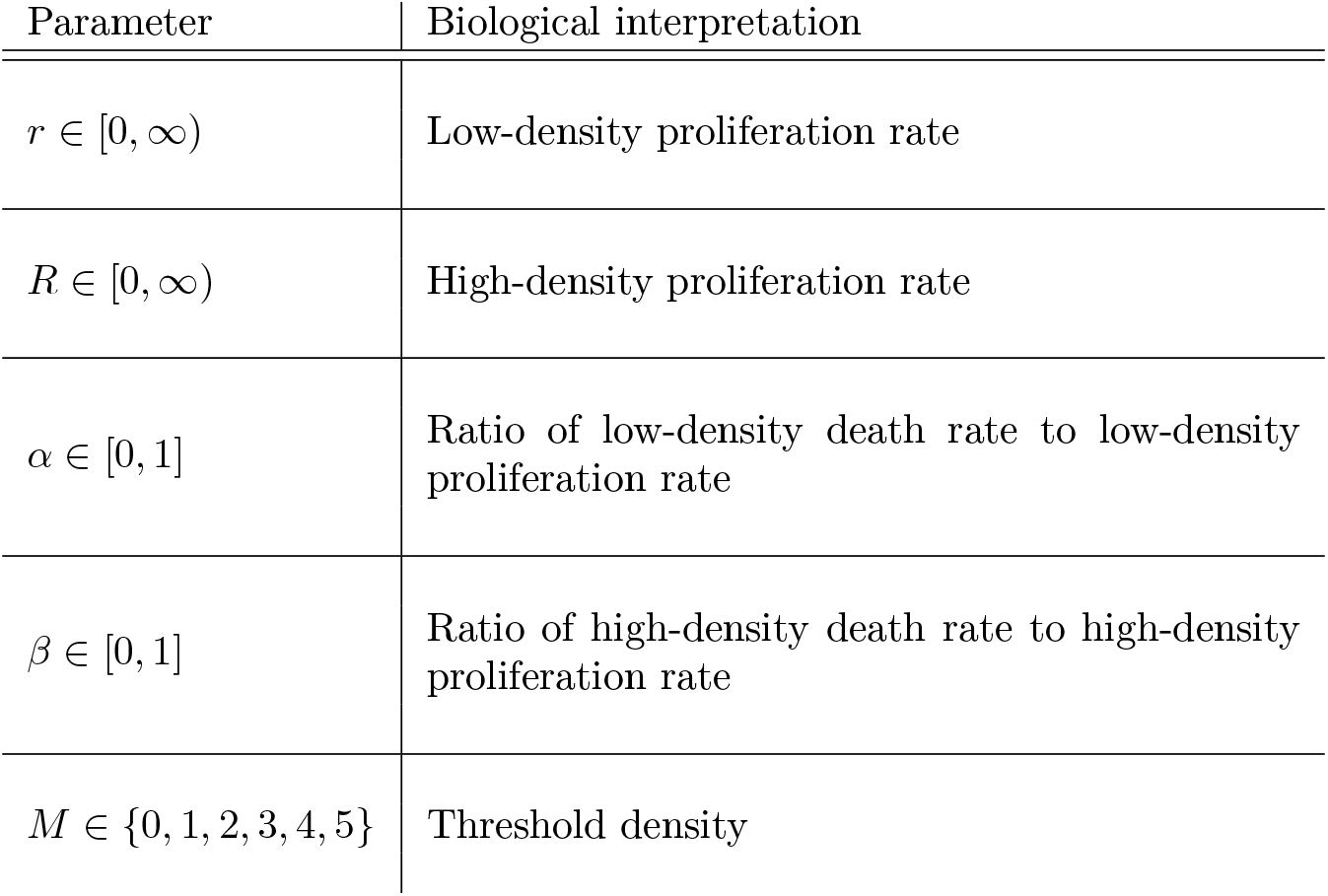
Summary of parameters used in the Binary Switch Model.

| Parameter | Biological interpretation |
| --- | --- |
| $r \in [0, \infty)$ | Low-density proliferation rate |
| $R \in [0, \infty)$ | High-density proliferation rate |
| $\alpha \in [0, 1]$ | Ratio of low-density death rate to low-density proliferation rate |
| $\beta \in [0, 1]$ | Ratio of high-density death rate to high-density proliferation rate |
| $M \in \{0, 1, 2, 3, 4, 5\}$ | Threshold density |

In Fig. 2, we show how the Binary Switch Model gives rise to non-logistic growth mechanisms. When no binary switch is present (Fig. 2a), the growth mechanisms are independent of local density and assume a single proliferation and death rate, resulting in logistic growth. However, when a binary switch is incorporated into the proliferation and death rates (Fig. 2b,c), the population dynamics described by (7) deviates from the classical logistic growth model. Consequently, we now wish to examine the various kinds of Allee effects the Binary Switch Model can give rise to. The main qualitative differences between logistic growth and various Allee effects are based on the number of equilibria and their stability; therefore, we now examine the roots of (7) for various parameter values. In all parameter regimes considered in the work, the zero equilibrium, *C*\* = 0, will always be present. Additional equilibria, if present, will be denoted as *C*\* = *C_i_* ∈ (0, 1], where *i* = 1, 2*, …* and are ordered such that *C_i_ < C_i_*_+1_ for all *i*. Since the right-hand side of (7) is a sixth-degree polynomial, a maximum of six equilibria can be present in (0, 1], but explicit expressions for the solutions of the polynomial cannot be determined in general. We will show that in the Binary Switch Model, a maximum of three equilibria can be present in (0, 1]. Setting *r* = 0 and *R >* 0 (Case 1) or *R* = 0 and *r >* 0 (Case 2), we will show that fewer equilibria are present in (0, 1]. In Case 3, corresponding to *r >* 0 and *R >* 0, certain combinations of parameter values produce equilibria with additional qualitative features, such as double-root and triple-root equilibria. For these special equilibria, we will designate particular symbols to *C_i_*, which appear as required.

### 2.1 Case 1: *r* = 0 and *R* > 0

This case corresponds to situations where individuals *below* the threshold density *M* do not proliferate or die. We will now show that in Case 1, either no equilibria are present in (0, 1], or we have one equilibrium *C*_1_ ∈ (0, 1], depending on the choice of *β* and *M*. In this regime, (7) simplifies to

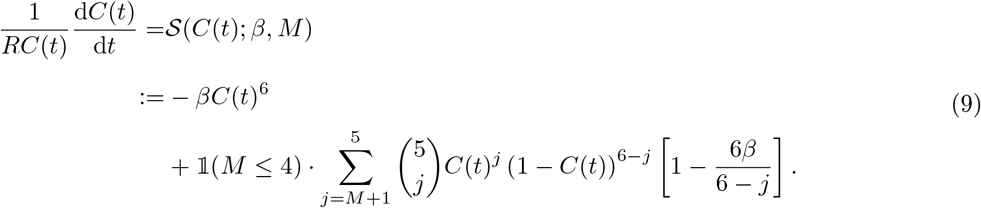

Since *β* appears as a linear coefficient in (9), it is easier to solve 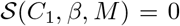 for *β* than for *C*_1_. The resulting relationship between *C*_1_ and *β* depends on the integer value of *M* ∈ {0, 1, 2, 3, 4, 5}; however, a general solution in terms of arbitrary *M* is difficult to obtain. Instead, we define the *family* of functions, *f_M_* (*C*_1_), for a particular value of *M*, such that

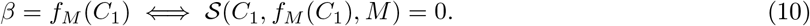

Using *f_M_* (*C*_1_), we determine the unique value of *β* that solves 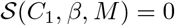 for a given value of *C*_1_ ∈ (0, 1], shown in Table 2. Plotting *β* = *f_M_* (*C*_1_) for all *M* ∈ {0, 1, 2, 3, 4, 5} and *C*_1_ ∈ (0, 1] indicates that *f_M_* (*C*_1_) is one-to-one on *C*_1_ ∈ (0, 1]. Therefore, the inverse function 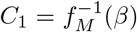 also has one solution, provided that *β* ∈ [0, (5 − *M*)/6). This range of *β* is obtained by mapping the *C*_1_ interval (0, 1] via the functions *f_M_* (*C*_1_). The functions *f_M_* (*C*_1_) in Table 2 provide a link between *β* and *C*_1_: if *C*_1_ is known, *β* = *f_M_* (*C*_1_) provides the parameter value to input in the model to obtain such an equilibrium. Conversely, if *β* is known, Table 2 indicates whether or not *C*_1_ ∈ (0, 1]. Finally, we note that when *β* ≥ (5 − *M*)/6, or when *M* = 5, only the zero equilibrium, *C*\* = 0, is present.

**Table 2:**
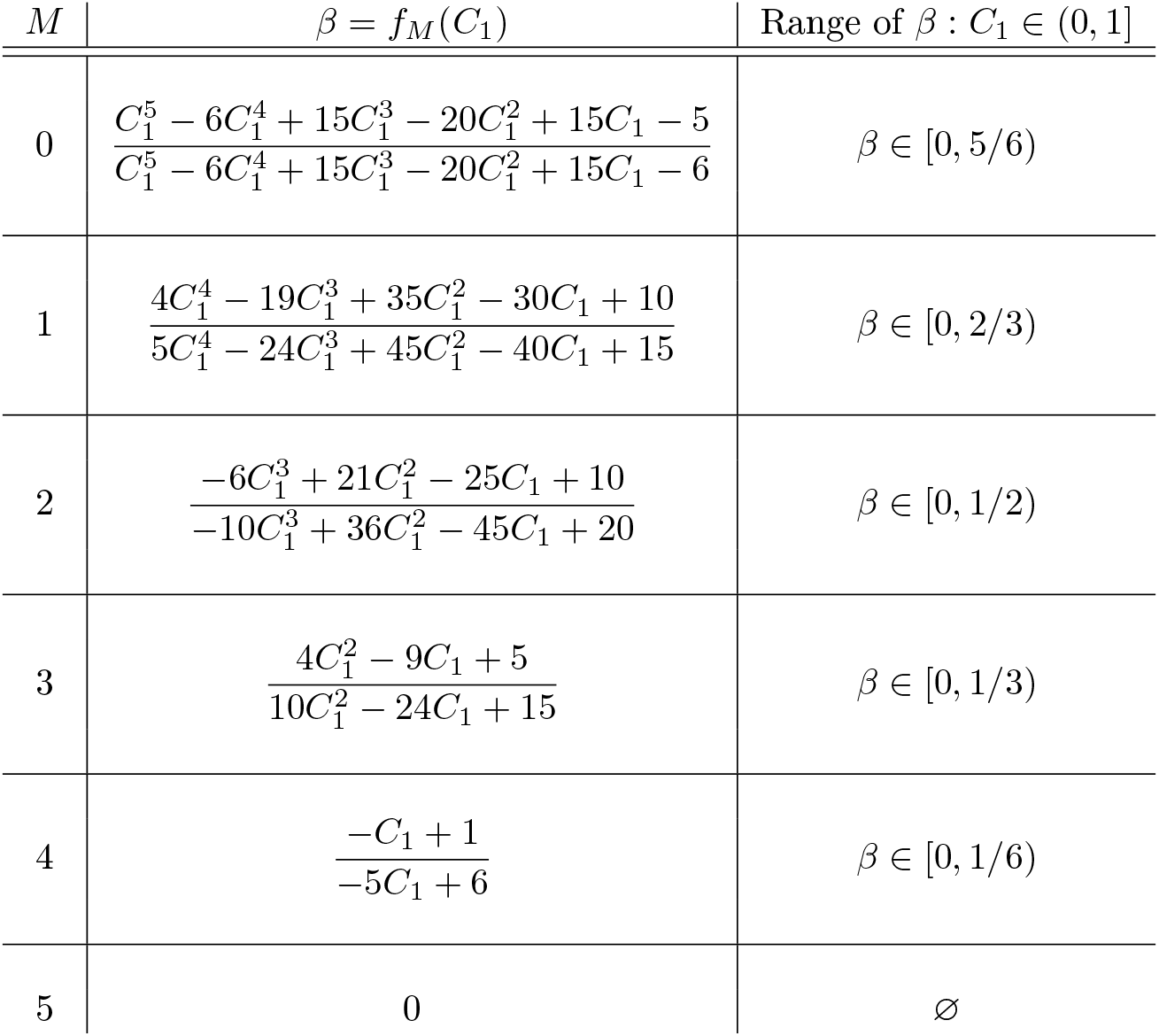
Relationships between the non-zero equilibrium of the Binary Switch Model, *C*_1_, to *β* and *M* for Case 1 when *r* = 0 (9).

| $M$ | $\beta = f_M(C_1)$ | Range of $\beta : C_1 \in (0, 1]$ |
| --- | --- | --- |
| 0 | $\frac{C_1^5 - 6C_1^4 + 15C_1^3 - 20C_1^2 + 15C_1 - 5}{C_1^5 - 6C_1^4 + 15C_1^3 - 20C_1^2 + 15C_1 - 6}$ | $\beta \in [0, 5/6)$ |
| 1 | $\frac{4C_1^4 - 19C_1^3 + 35C_1^2 - 30C_1 + 10}{5C_1^4 - 24C_1^3 + 45C_1^2 - 40C_1 + 15}$ | $\beta \in [0, 2/3)$ |
| 2 | $\frac{-6C_1^3 + 21C_1^2 - 25C_1 + 10}{-10C_1^3 + 36C_1^2 - 45C_1 + 20}$ | $\beta \in [0, 1/2)$ |
| 3 | $\frac{4C_1^2 - 9C_1 + 5}{10C_1^2 - 24C_1 + 15}$ | $\beta \in [0, 1/3)$ |
| 4 | $\frac{-C_1 + 1}{-5C_1 + 6}$ | $\beta \in [0, 1/6)$ |
| 5 | 0 | ∅ |

To determine the stability of the equilibria, we consider the cases when *β* ∈ [0, (5 − *M*)/6) and when *β* ≥ (5 − *M*)/6 separately. When *β* ∈ [0, (5 − *M*)/6), two distinct equilibria are present: *C*\* = 0 and *C*\* = *C*_1_ ∈ (0, 1]. Based on the sign of 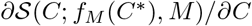 at these equilibria, *C*\* = 0 is always unstable and *C*\* = *C*_1_ is always stable. These features are consistent with the *Weak Allee effect*, whereby the density growth rate deviates from logistic growth without incorporating additional equilibria. When *β* ≥ (5 − *M*)/6, or when *M* = 5, *C*\* = 0 is the only equilibrium and it is always stable, corresponding to the qualitative features of an *extinction* density growth rate, where 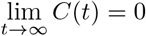 for all *C*(0). Both qualitative features in this parameter regime are shown in the bifurcation diagram in Fig. 3. We conclude that in Case 1, either zero or one equilibria is present in the interval (0, 1], corresponding to extinction and Weak Allee parameter regimes, respectively.

**Figure 3:**
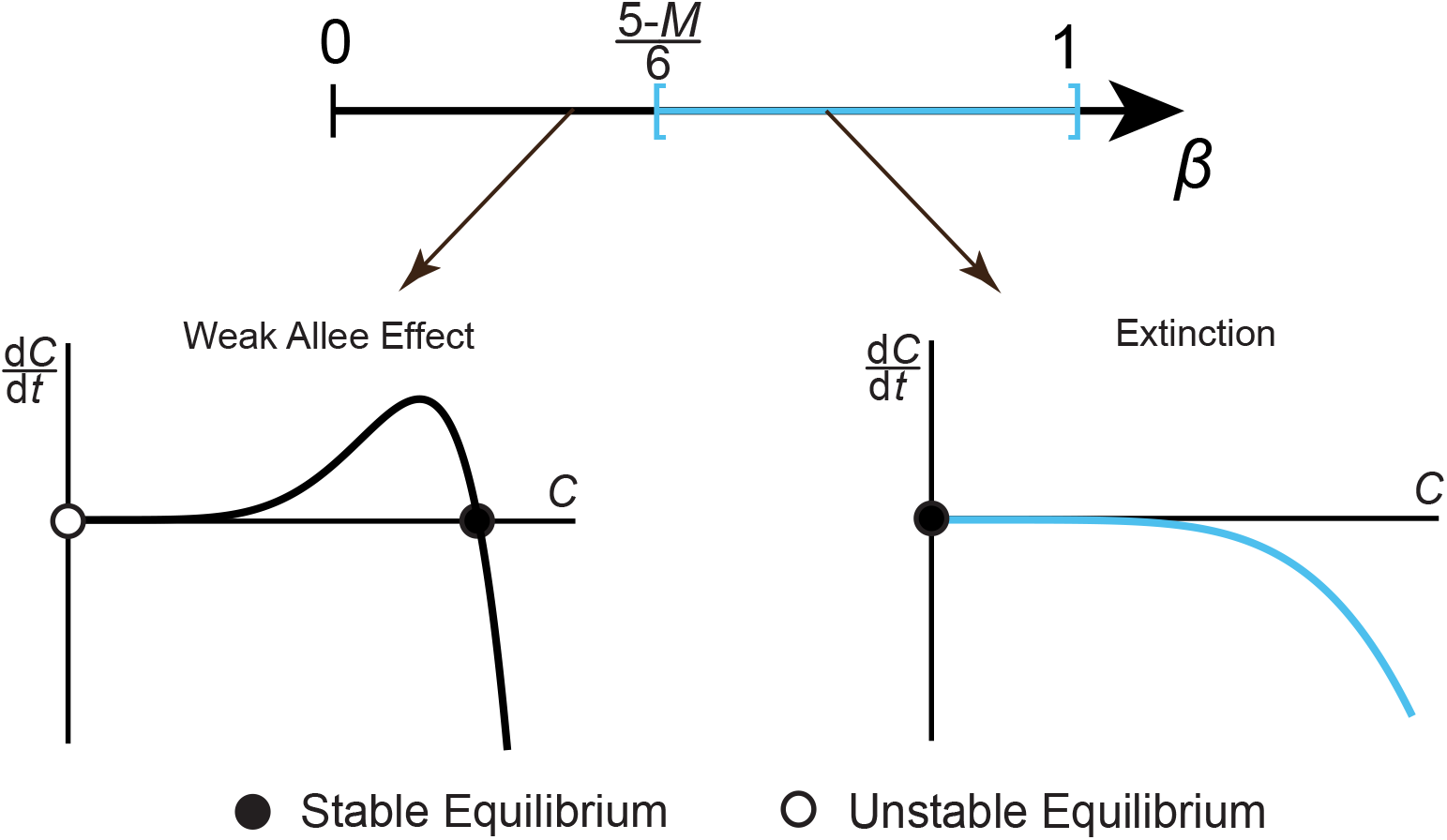
Bifurcation diagram of the Binary Switch Model, shown in (9), for Case 1 when *r* = 0. Varying *β* produces different qualitative features in terms of equilibria and their stability. The resulting density growth rates, d*C/*d*t*, are shown as a function of *C*, where a stable equilibrium is represented with a black circle and an unstable equilibrium with a white circle.

### 2.2 Case 2: *r* > 0 and *R* = 0

This case corresponds to when individuals *above M* do not proliferate or die. When *R* = 0, we have

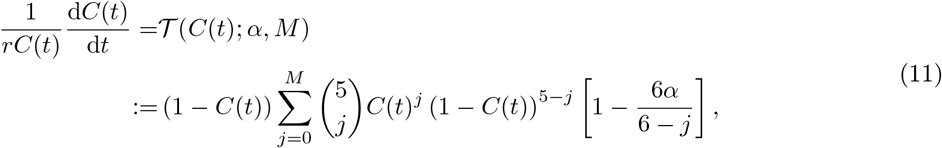

which is independent of *β*. In a similar fashion to Case 1, we consider the equilibria for various choices of *α* and *M*, noting that *C*\* = 0 and *C*\* = 1 are always equilibria in this case. However, we will show that in Case 2, we have the possibility of a third equilibrium in (0, 1). When this additional equilibria is present, then *C*_2_ = 1 and *C*_1_ ∈ (0, 1); otherwise, *C*_1_ = 1. To determine if *C*\* = 1 is the first or second non-zero equilibrium, we define

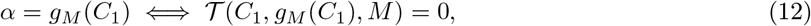

and determine the value of *α* that solves 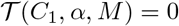 for a given value of *C*_1_ ∈ (0, 1), shown in Table 3. Like Case 1, the family of functions *α* = *g_M_* (*C*_1_) provide an explicit relationship between *α* and *C*_1_. Since *α* = *g_M_* (*C*_1_) is one-to-one on *C*_1_ ∈ (0, 1), the inverse function 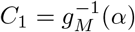 also has one solution, *C*_1_ ∈ (0, 1), provided *α* ∈ ((6 − *M*)/6, 1). This value of *C*_1_ ∈ (0, 1) provides a *third* equilibrium of (11); conversely, when *α* ≤ (6 − *M*)/6, or when *M* = 0, the only two equilibria are *C*\* = 0 and *C*_1_ = 1.

**Table 3:** Relation between non-zero equilibrium, 0 ≤ *C*_1_ < 1, to *α* and *M* for Case 2 when *R* = 0 (11).

| $M$ | $\alpha = g_M(C_1)$ | Range of $\alpha : C_1 \in (0, 1)$ |
| --- | --- | --- |
| 0 | 1 | ∅ |
| 1 | $\frac{4C_1 + 1}{5C_1 + 1}$ | $\alpha \in (5/6, 1)$ |
| 2 | $\frac{6C_1^2 + 3C_1 + 1}{10C_1^2 + 4C_1 + 1}$ | $\alpha \in (2/3, 1)$ |
| 3 | $\frac{4C_1^3 + 3C_1^2 + 2C_1 + 1}{10C_1^3 + 6C_1^2 + 3C_1 + 1}$ | $\alpha \in (1/2, 1)$ |
| 4 | $\frac{C_1^4 + C_1^3 + C_1^2 + C_1 + 1}{5C_1^4 + 4C_1^3 + 3C_1^2 + 2C_1 + 1}$ | $\alpha \in (1/3, 1)$ |
| 5 | $\frac{1}{C_1^5 + C_1^4 + C_1^3 + C_1^2 + C_1 + 1}$ | $\alpha \in (1/6, 1)$ |

In the case where *C*_1_ ∈ (0, 1), examining the sign of 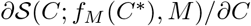 shows that *C*\* = 0 and *C*\* = 1 are unstable, whereas *C*\* = *C*_1_ is stable. This combination of equilibria has the opposite stability properties of the Strong Allee effect (3), and so we refer to density growth rates with these stability properties as the *Reverse Allee effect*. In the case where *α* ≤ (6 − *M*)/6, or when *M* = 0, stability analysis shows that *C*_1_ = 1 is stable and *C*\* = 0 is unstable, which is consistent with the qualitative features of the Weak Allee effect. Finally, when *α* = 1, we return to having only two equilibria, *C*\* = 0 and *C*\* = 1, but the stability is the opposite of the usual Weak Allee effect. Therefore, when *α* = 1, 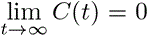 for *C*(0) < 1. All these qualitative features in this parameter regime are shown in the bifurcation diagram in Fig. 4. We conclude that in Case 2, either one or two equilibria are present in (0, 1], with the *Extinction* regime occurring when *α* = 1. For *α <* 1, a new kind of Allee effect, which we call the *Reverse Allee effect*, occurs if two equilibria are present in (0, 1]; otherwise, we retrieve the Weak Allee effect.

**Figure 4:**
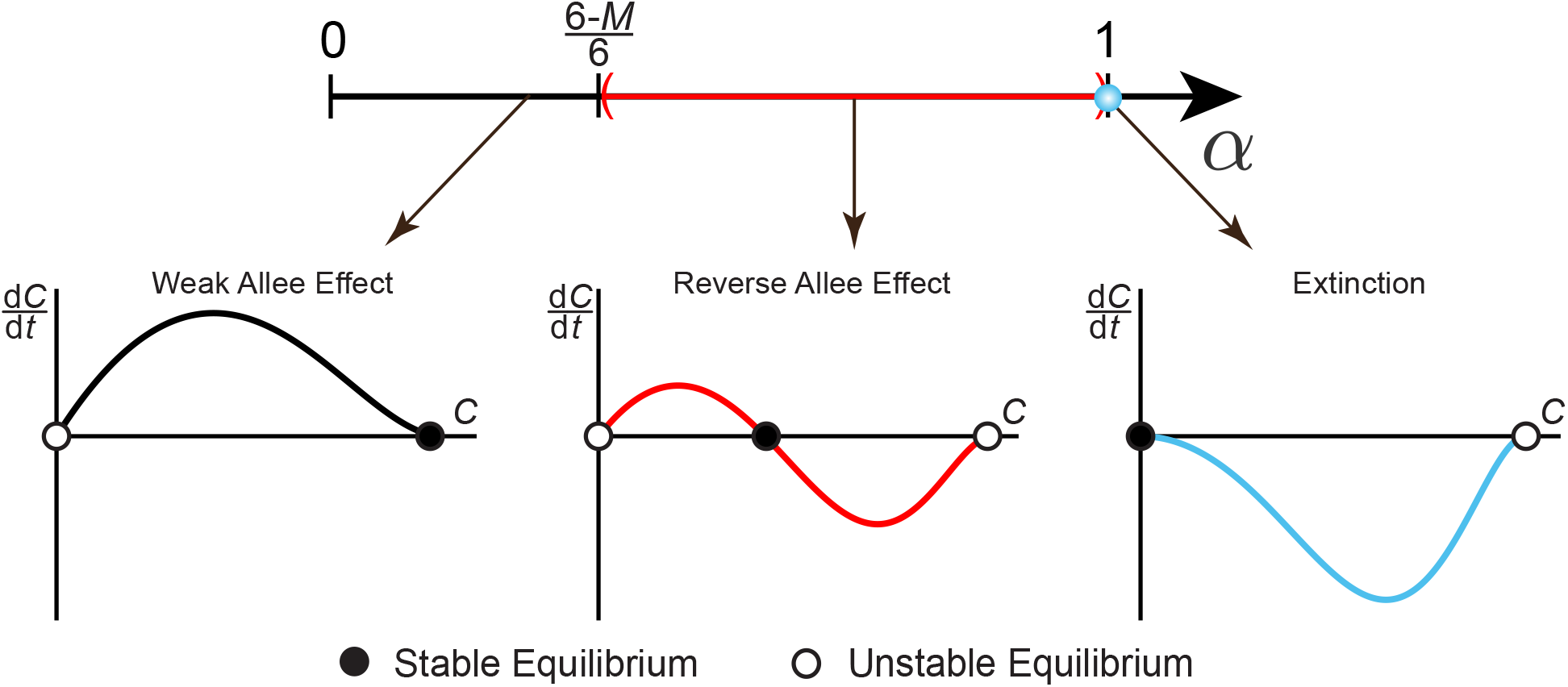
Bifurcation diagram of the Binary Switch Model, shown in (11), for Case 2 when *R* = 0. Varying *α* produces different qualitative features in terms of equilibria and their stability. The resulting density growth rates, d*C/*d*t*, are shown as a function of *C*, where a stable equilibrium is represented with a black circle and an unstable equilibrium with a white circle.

### 2.3 Case 3: *r* > 0 and *R* > 0

In the most general case, the proliferation and death rates of individuals change at the threshold density *M*, but remain non-zero on either side of the threshold density. As a result, (7) can be written as

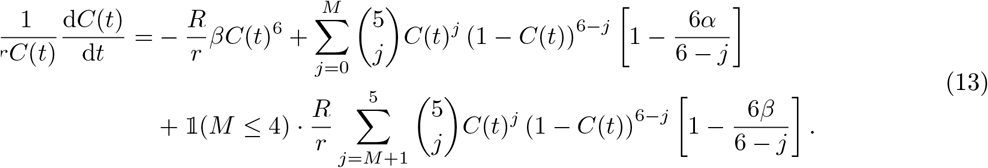

Without loss of generality, we assume that *r* = 1, since other non-zero values or *r* can be rescaled to unity by changing the timescale in (7), which does not affect its equilibria. Consequently, with some rearranging, we have

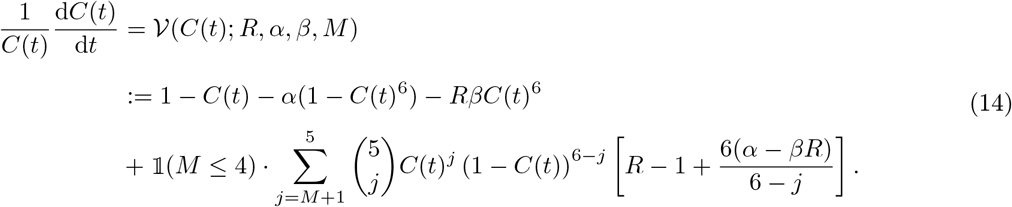

We will show that in Case 3, there can be between zero and three equilibria in (0, 1], noting that *C*\* = 1 is an equilibrium of (14) if and only if *β* = 0. When we have three distinct equilibria in (0, 1], we obtain a new type of Allee effect, referred to here as the *Hyper-Allee effect* (Fadai et al., 2019), in which the zero equilibrium is unstable, and an intermediate unstable equilibrium is contained between two positive, stable equilibria. However, in order for the parameter space to continuously transition from the Weak Allee effect, as in Cases 1 and 2, to the Hyper-Allee effect, there must exist a critical set of model parameters at which a *double-root* equilibrium occurs. Therefore, to determine what regions of (*R, α, β, M*) parameter space exhibit Hyper-Allee effects instead of the Weak Allee effect, we focus on determining the *boundary* of these effects in terms of model parameters and equilibria. This boundary, defined as the *Tangential Manifold*, will be the focus of our analysis in this section.

In addition to determining the boundary between Weak Allee and Hyper-Allee parameter spaces, we will also show that even more Allee effects are present when *α* = 1. In particular, we show that in Case 3, the Extinction parameter regime continues to exist, along with the Strong Allee effect, when *α* = 1. We also determine an explicit relationship between *R, β,* and *M* for when the Extinction regime becomes the Strong Allee effect, which is linked to the Tangential Manifold. We now focus our attention on determining additional equilibria *C_i_* ∈ (0, 1].

Numerical observations indicate that certain combinations of (*R, α, β, M*) can produce up to three distinct values of *C_i_* ∈ (0, 1] satisfying 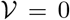. Furthermore, in parameter regimes where three distinct equilibria are present in (0, 1], stability analysis about these equilibria reveals that *C*\* = 0 and *C*\* = *C*_2_ are unstable equilibria, whereas *C*\* = *C*_1_ and *C*\* = *C*_3_ are stable equilibria. These qualitative features are consistent with the aforementioned Hyper-Allee effect, which is a higher-order effect that is very different to the usual Weak Allee and Strong Allee effects (Fig. 5).

**Figure 5:**
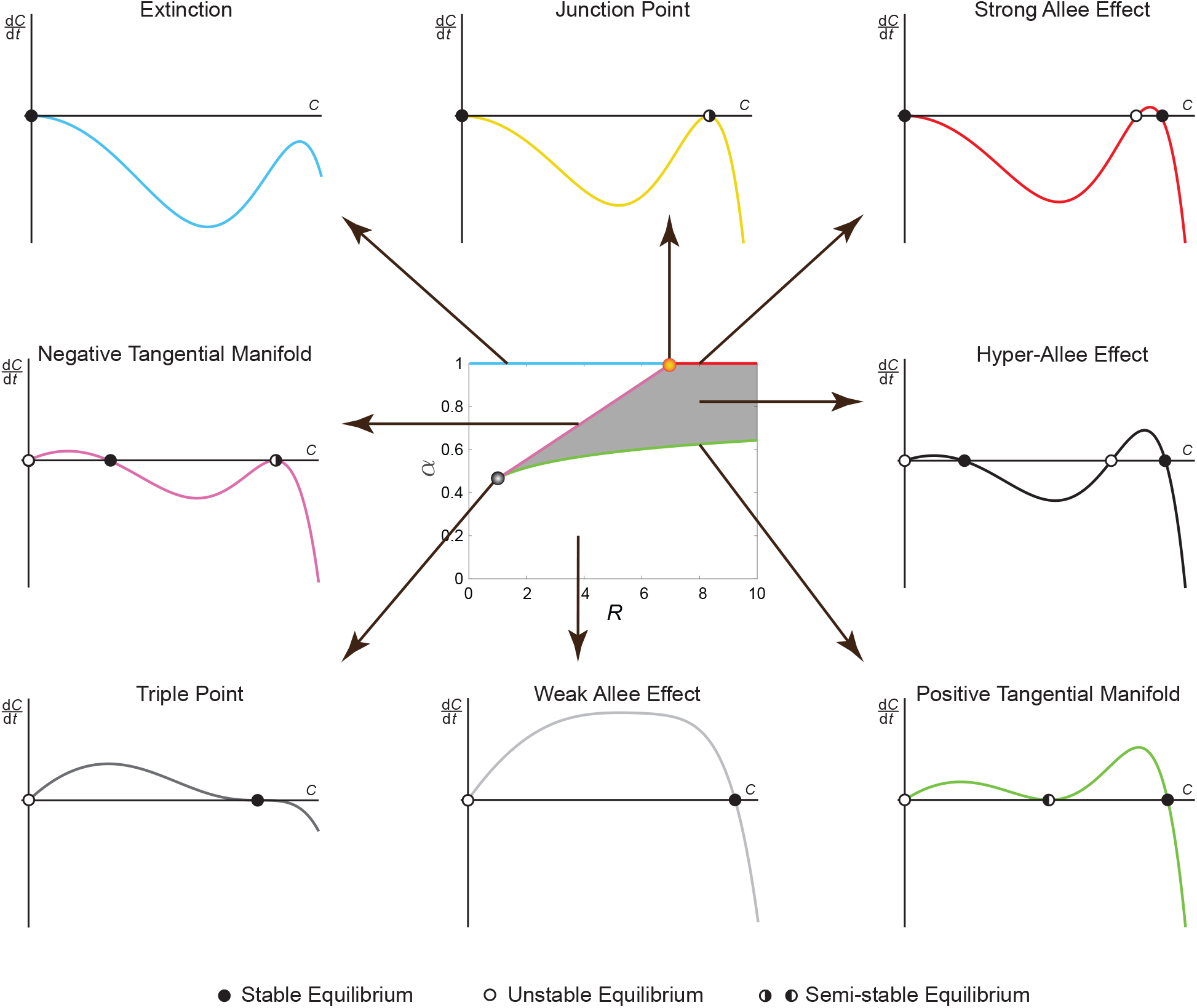
Bifurcation diagram of the Binary Switch Model for Case 3, shown in (14), with *β* = 0.06*, r* = 1*, R >* 0, and *M* = 4. Pairs of (*α, R*) parameters produce different qualitative features, in terms of equilibria and their stability. The resulting density growth rates, d*C/*d*t*, are shown as a function of *C*, where a stable equilibrium is represented with a black circle, an unstable equilibrium with a white circle, and a semi-stable equilibrium with a half-filled circle.

For solutions to continuously transition from one equilibrium in (0, 1], like the Weak Allee effect in Cases 1 and 2, to three equilibria in (0, 1], such as the Hyper-Allee effect, we must have certain values of (*R, α, β, M*) that produce a *double root* for *C_i_*. We denote this special case of a double root equilibrium as 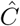, which can occur in either the *C*_1_ or *C*_2_ equilibrium position. In addition to satisfying 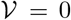, the double root equilibrium, 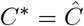, must also satisfy

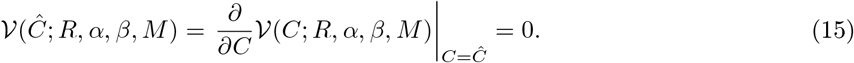

The set of parameters satisfying (15) is referred to as the *Tangential Manifold*, where the double root equilibrium, 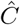, is a *semi-stable equilibrium* of (14) (Strogatz, 2018). A semi-stable equilibrium 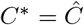 has the properties that populations slightly larger than 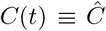 remain close to 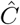, but populations slightly smaller than 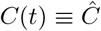 diverge away from 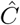, or vice-versa. Since we have two equations with four unknowns, we parametrise the Tangential Manifold as 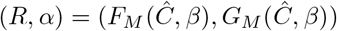, for particular values of 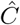 and *β* (Fig. 5). The functions 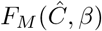 and 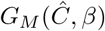 describing the Tangential Manifold are shown in Table 4.

**Table 4:** Relation between the semi-stable equilibrium, 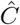, to *α, β, R* and *M* for Case 3. Parameter values satisfying 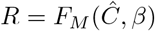 and 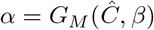 are members of the Tangential Manifold. If 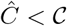, then 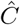 is a member of the Positive Tangential Manifold; if 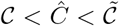, then 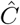 is a member of the Negative Tangential Manifold. The Triple Point, 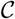, is defined implicitly via 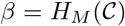, while the Junction Point, 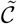, is defined implicitly via 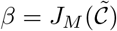.

| $M$ | $R = F_M(\hat{C}, \beta)$ | $\beta = H_M(\mathcal{C})$ |
| --- | --- | --- |
| 0 | 0 | ∅ |
| 1 | $\frac{(\hat{C} - 1)^6}{\hat{C}(\hat{C}^5 - 6\hat{C}^4 + 15\hat{C}^3 - 20\hat{C}^2 - 10\hat{C} - 30\beta + 20)}$ | $\frac{2(1 - \mathcal{C})}{3}$ |
| 2 | $\frac{(\hat{C} - 1)^5(6\hat{C}^2 + 8\hat{C} + 1)}{\hat{C}^2(6\hat{C}^5 - 22\hat{C}^4 + 21\hat{C}^3 + 15\hat{C}^2 + 10\hat{C} + 60\beta - 30)}$ | $\frac{(1 - \mathcal{C})(1 + 2\mathcal{C})}{3\mathcal{C} + 2}$ |
| 3 | $\frac{(\hat{C} - 1)^4(6\hat{C}^4 + 16\hat{C}^3 + 21\hat{C}^2 + 6\hat{C} + 1)}{\hat{C}^3(6\hat{C}^5 - 8\hat{C}^4 - 7\hat{C}^3 - 6\hat{C}^2 - 5\hat{C} - 60\beta + 20)}$ | $\frac{(1 - \mathcal{C})(1 + 2\mathcal{C} + 2\mathcal{C}^2)}{3\mathcal{C}^2 + 4\mathcal{C} + 3}$ |
| 4 | $\frac{(\hat{C} - 1)^3(\hat{C}^6 + 4\hat{C}^5 + 10\hat{C}^4 + 20\hat{C}^3 + 10\hat{C}^2 + 4\hat{C} + 1)}{\hat{C}^4(\hat{C}^5 + \hat{C}^4 + \hat{C}^3 + \hat{C}^2 + \hat{C} + 30\beta - 5)}$ | $\frac{(1 - \mathcal{C}^2)(2\mathcal{C}^2 + \mathcal{C} + 2)}{3(\mathcal{C}^3 + 2\mathcal{C}^2 + 3\mathcal{C} + 4)}$ |
| 5 | 0 | ∅ |
| $M$ | $\alpha = G_M(\hat{C}, \beta)$ | $\beta = J_M(\tilde{\mathcal{C}})$ |
| 0 | 1 | ∅ |
| 1 | $\frac{\beta(\hat{C}^5 - 6\hat{C}^4 + 15\hat{C}^3 - 20\hat{C}^2 + 15\hat{C} - 30) - 20(\hat{C} - 1)}{\hat{C}^5 - 6\hat{C}^4 + 15\hat{C}^3 - 20\hat{C}^2 - 10\hat{C} - 30\beta + 20}$ | $\frac{(\tilde{\mathcal{C}} - 1)(\tilde{\mathcal{C}}^3 - 5\tilde{\mathcal{C}}^2 + 10\tilde{\mathcal{C}} - 10)}{\tilde{\mathcal{C}}^4 - 6\tilde{\mathcal{C}}^3 + 15\tilde{\mathcal{C}}^2 - 20\tilde{\mathcal{C}} + 15}$ |
| 2 | $\frac{\beta(6\hat{C}^5 - 22\hat{C}^4 + 21\hat{C}^3 + 15\hat{C}^2 - 40\hat{C} + 60) + 30(\hat{C} - 1)}{6\hat{C}^5 - 22\hat{C}^4 + 21\hat{C}^3 + 15\hat{C}^2 + 10\hat{C} + 60\beta - 30}$ | $\frac{(\tilde{\mathcal{C}} - 1)(6\tilde{\mathcal{C}}^3 - 16\tilde{\mathcal{C}}^2 + 5\tilde{\mathcal{C}} + 20)}{6\tilde{\mathcal{C}}^4 - 22\tilde{\mathcal{C}}^3 + 21\tilde{\mathcal{C}}^2 + 15\tilde{\mathcal{C}} - 40}$ |
| 3 | $\frac{\beta(6\hat{C}^5 - 8\hat{C}^4 - 7\hat{C}^3 - 6\hat{C}^2 + 45\hat{C} - 60) - 20(\hat{C} - 1)}{6\hat{C}^5 - 8\hat{C}^4 - 7\hat{C}^3 - 6\hat{C}^2 - 5\hat{C} - 60\beta + 20}$ | $\frac{(\tilde{\mathcal{C}} - 1)(6\tilde{\mathcal{C}}^3 - 2\tilde{\mathcal{C}}^2 - 9\tilde{\mathcal{C}} - 15)}{6\tilde{\mathcal{C}}^4 - 8\tilde{\mathcal{C}}^3 - 7\tilde{\mathcal{C}}^2 - 6\tilde{\mathcal{C}} + 45}$ |
| 4 | $\frac{\beta(\hat{C}^5 + \hat{C}^4 + \hat{C}^3 + \hat{C}^2 - 24\hat{C} + 30) + 5(\hat{C} - 1)}{\hat{C}^5 + \hat{C}^4 + \hat{C}^3 + \hat{C}^2 + \hat{C} + 30\beta - 5}$ | $\frac{(\tilde{\mathcal{C}} - 1)(\tilde{\mathcal{C}}^3 + 2\tilde{\mathcal{C}}^2 + 3\tilde{\mathcal{C}} + 4)}{\tilde{\mathcal{C}}^4 + \tilde{\mathcal{C}}^3 + \tilde{\mathcal{C}}^2 + \tilde{\mathcal{C}} - 24}$ |
| 5 | 1/6 | ∅ |

While the Tangential Manifold can be determined explicitly by solving (15), we observe that two forms of a semi-stable equilibrium can occur (Fig. 5). If the double root 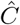 is below some critical value, 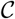, then this semi-stable equilibrium occurs between *C*\* = 0, which is unstable, and some larger equilibrium *C*\* = *C*_2_, which is stable. If 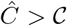, then this semi-stable equilibrium is larger than both *C*\* = 0 and *C*\* = *C*_1_, which remain unstable and stable, respectively. We refer to the branch of the Tangential Manifold where 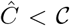 as the *Positive Tangential Manifold*, based on the sign of the density growth rate between 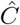 and *C*_2_ (Fig. 5). In a similar fashion, we refer to the branch of the Tangential Manifold where 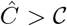 as the *Negative Tangential Manifold*. When 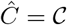, the double root becomes a stable triple root and 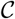 satisfies

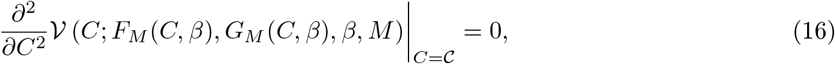

where 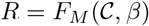 and 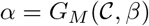 are chosen to ensure we remain on the Tangential Manifold. Equation (16) provides an additional constraint on the Tangential Manifold, implying that we can relate 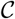 to a unique value of *β*. We denote 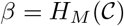 if (16) is satisfied, with 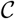 denoting the *Triple Point* of (14) (Table 4).

Additionally, from Fig. 5, we note that when *α* = 1, the equilibria and their resulting stability change, compared to *α <* 1. When *α* = 1, the Negative Tangential Manifold is valid for a unique pair of (*β, R*) parameters, for a particular equilibrium value, 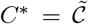. We define this critical equilibrium value as the *Junction Point*, which satisfies

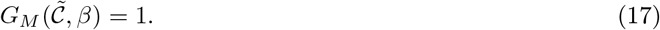

We denote 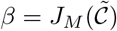 if (17) is satisfied (Table 4); furthermore, we determine the corresponding value of *R* at the Junction Point by evaluating 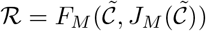. When *α* = 1 and 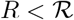, the only equilibrium value of (14) is *C*\* = 0, which is stable. This implies that all population densities go extinct in this parameter regime. When *α* = 1 and 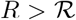, (14) has three solutions: *C*\* = 0, which is stable, an intermediate-valued unstable equilibrium *C*\* = *C*_1_, and a larger-valued stable equilibrium *C*\* = *C*_2_ (Fig. 5). Thus, the stability features of this density growth rate are the same as the *Strong Allee effect*. When 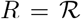, the Junction Point, 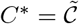, is semi-stable, while *C*\* = 0 remains stable. A summary of this diverse family of Allee effects, in terms of the number and stability of the equilibria, is shown in Table 5.

**Table 5:** Summary of qualitative features seen in the Binary Switch Model. The stability of each equilibrium, listed in increasing order of magnitude, can be stable (*S*), unstable (*U*), or semi-stable (*SS*).

| Effect name | Equilibria | Stability | Notes |
| --- | --- | --- | --- |
| Extinction | $\{0\}$ | $\{S\}$ | |
| Logistic Growth | $\{0, C_1\}$ | $\{U, S\}$ | $r = R, \alpha = \beta$ |
| Weak Allee / Triple Point | $\{0, C_1\}$ | $\{U, S\}$ | Triple: $C_1 = \mathcal{C}$ |
| Junction Point | $\{0, C_1\}$ | $\{S, SS\}$ | $C_1 = \tilde{\mathcal{C}}$ |
| Strong Allee | $\{0, C_1, C_2\}$ | $\{S, U, S\}$ | |
| Reverse Allee | $\{0, C_1, C_2\}$ | $\{U, S, U\}$ | $C_2 = 1$ |
| Positive Tangential Manifold | $\{0, C_1, C_2\}$ | $\{U, SS, S\}$ | $C_1 = \hat{\mathcal{C}}$ |
| Negative Tangential Manifold | $\{0, C_1, C_2\}$ | $\{U, S, SS\}$ | $C_2 = \hat{\mathcal{C}}$ |
| Hyper-Allee | $\{0, C_1, C_2, C_3\}$ | $\{U, S, U, S\}$ | |

From Table 4, we note some key features of the Tangential Manifold. Firstly, when *β* = 0, we note that the Triple Point is 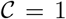 for 1 ≤ *M* ≤ 4. Since the Negative Tangential Manifold must have 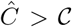, we conclude that the Negative Tangential Manifold does not exist when *β* = 0, which is also observed in Fig. 6. When *β* = (5 − *M*)/6 and 1 ≤ *M* ≤ 4, the Triple Point *and* the Junction Point are both 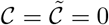, implying that no points are contained in the Tangential Manifold. Consequently, parameter pairs (*α, R*) that result in qualitative features other than the Extinction regime or the Weak Allee effect can only occur when *α <* 1 and *β* ∈ [0, (5 − *M*)/6), as shown in Fig. 6. Finally, we note that when *M* = 0 or *M* = 5, the Tangential Manifold does not exist, since the solution of (15) requires *R* = 0. Therefore, the qualitative features of (14) in the entire (*α, R*) parameter space are those seen in the Weak Allee effect when *α <* 1 and the Extinction regime when *α* = 1.

**Figure 6:**
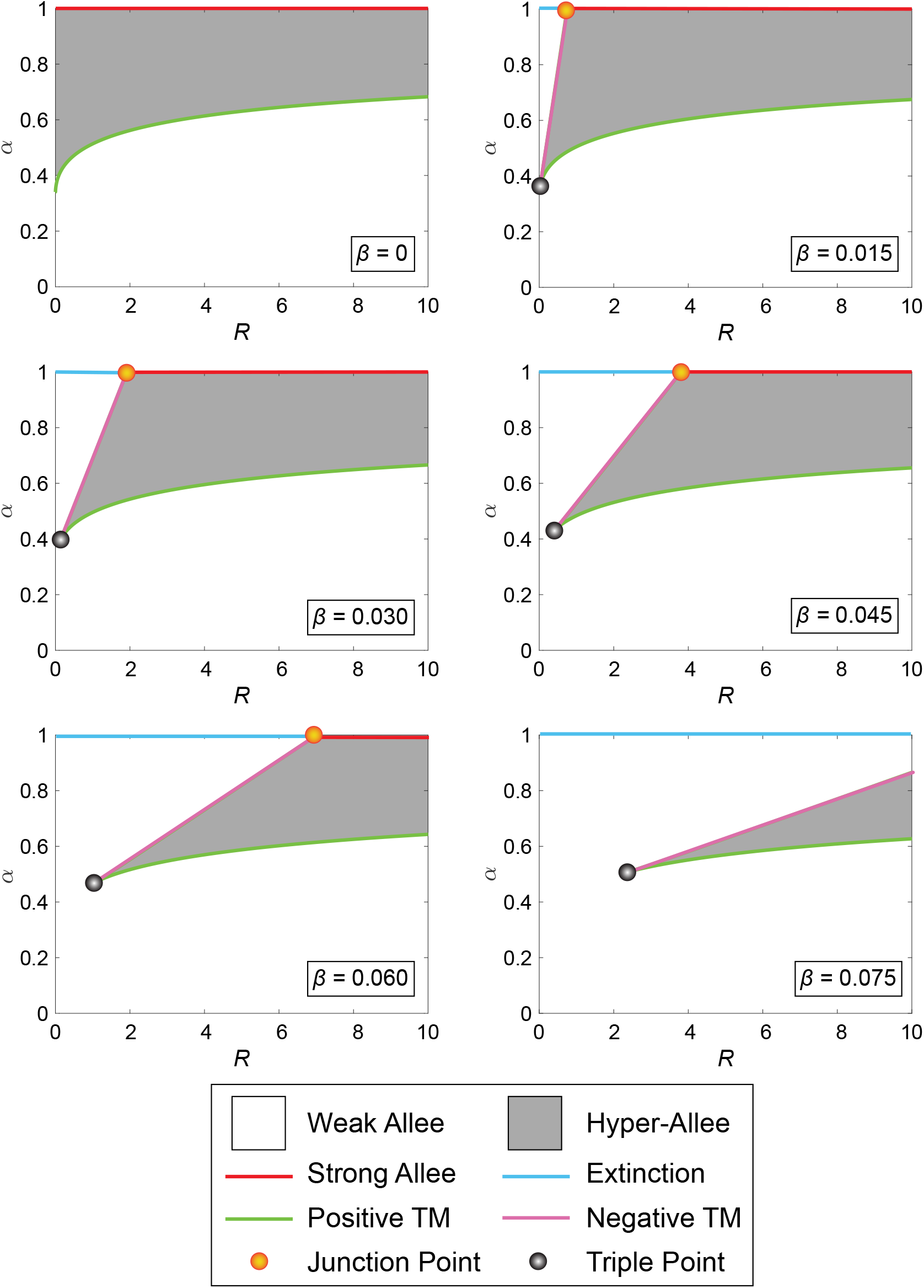
Bifurcation diagram of the Binary Switch Model for Case 3, shown in (14), with *M* = 4*, r* = 1*, R >* 0, and varying *β*. The qualitative forms of various effects are shown in the legend, described in further detail in Fig. 5. The parameter space exhibiting Hyper-Allee features vanishes as *β* → 1/6.

To summarise, we determine that in Case 3 when *M* ∈ {1, 2, 3, 4}, and *β* ∈ [0, (5−*M*)/6), a diverse family of Allee effects can be found. Among these Allee effects are: the Weak Allee effect, the Extinction regime, the Strong Allee effect, and a *Hyper-Allee effect* parameter regime with three distinct equilibria in (0, 1]. Additional Allee effects can be observed at the boundaries of the aforementioned Allee effects, including the Tangential Manifold and Junction Point with semi-stable equilibria, as well as the Triple Point with a single stable equilibria in (0, 1]. In all of these cases, there are between zero and three equilibria in the interval (0, 1].

## 3 Interpreting experimental data using the Binary Switch Model

To demonstrate how the Binary Switch Model can be used to provide biological insight, we consider population-level datasets describing the growth of populations of cancer cells. Neufeld et al. (2017) perform three experiments with U87 glioblastoma cells. Uniform monolayers of cells are grown from three different initial densities, with the data shown in Fig. 7. Here, we see that all three experiments lead to increasing population densities with time. The two experiments with the smallest initial densities lead to increasing, concave up *C*(*t*) profiles. The experiment with the largest initial density leads to an increasing *C*(*t*) profile that changes concavity at approximately *t* = 100 h.

**Figure 7:**
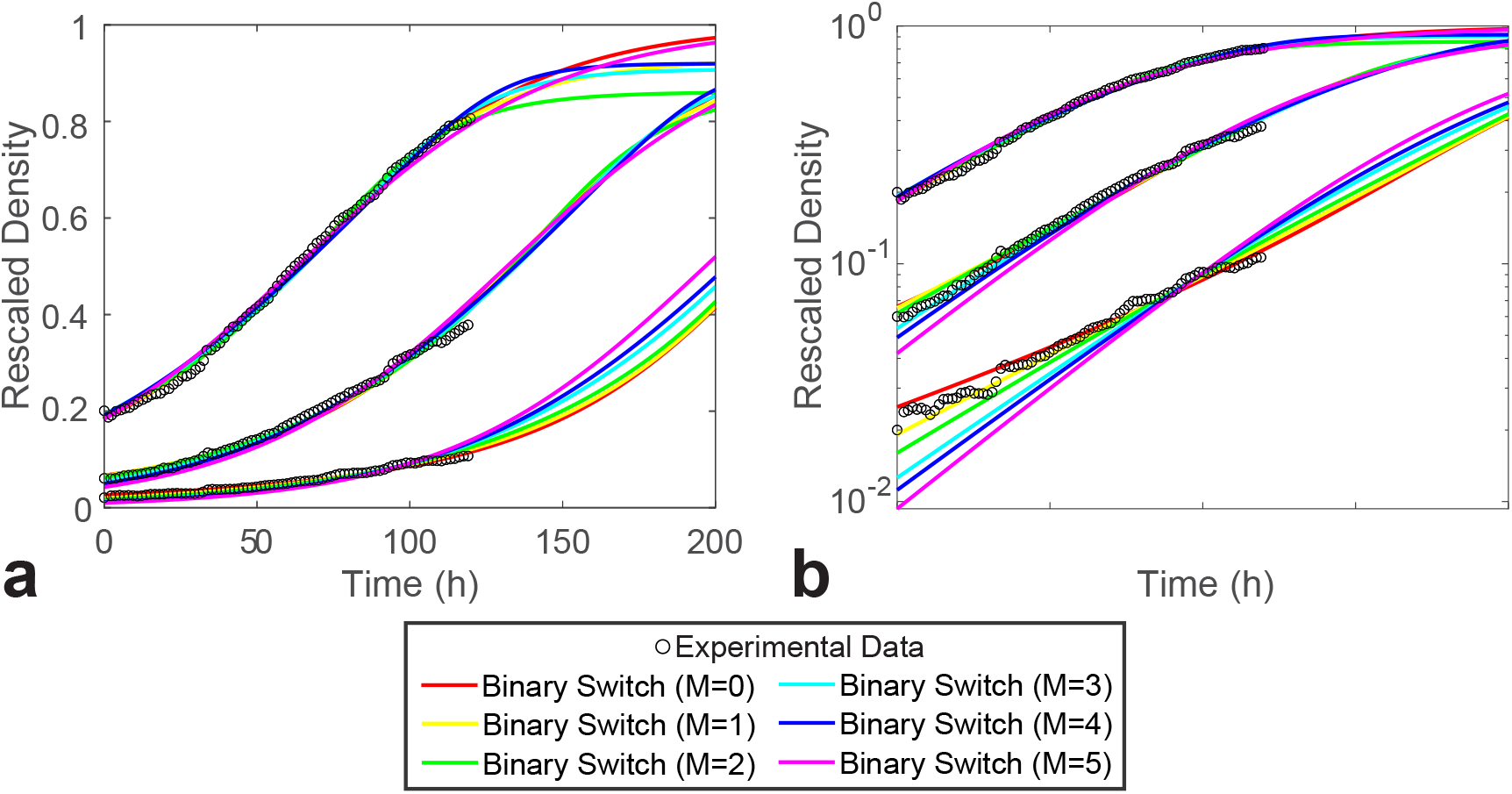
Population density of U87 glioblastoma cells compared to the calibrated Binary Switch Model. U87 glioblastoma cells, with initial densities of *c*_1_(0) = 0.02, *c*_2_(0) = 0.06, and *c*_3_(0) = 0.2, are observed over the span of 120 hours (black circles) (Neufeld et al., 2017). The Binary Switch Model (solid curves) is fit to minimise the combined least-square error (18), Σ*χ*^2^, of three experimental datasets shown in Neufeld et al. (2017). The estimates of the optimal model parameter set, for each value of *M*, is shown in Table 6. (b) A semi-log plot makes it easier to visually compare the quality of match between the data and the model.

The density of U87 glioblastoma cells has already been rescaled by its maximum packing density in Neufeld et al. (2017), so we assume that *C* = 1 corresponds to the maximum rescaled density. Our aim is to choose **Θ** = (*α, β, r, R, M*), with *C*_1_(0)*, C*_2_(0), and *C*_3_(0) as initial conditions, such that the model parameters provide the best match to all three experimental conditions simultaneously. It is important to calibrate the model to match all three datasets simultaneously, because if (7) is consistent with the experimental data, there should be a single choice of model parameters that matches the observed population dynamics, regardless of initial density (Jin et al., 2016b).

To match all experimental datasets simultaneously, we consider the combined least-squares error between model predictions and all data:

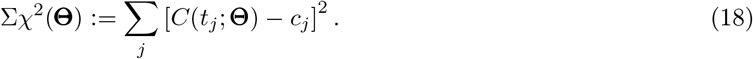

Here, we treat the initial densities, *C*_1_(0)*, C*_2_(0)*, C*_3_(0) as parameters to be determined; therefore we consider the extended parameter vector, **Θ** = (*M, r, R, α, β, C*_1_(0)*, C*_2_(0)*, C*_3_(0)). In (18), *c_j_* represents all three experimental datasets obtained at times *t_j_*, concatenated into a single vector. While the Binary Switch Model uses the initial conditions *C*_1_(0)*, C*_2_(0), and *C*_3_(0), we denote the experimental measurements at *t* = 0 h as *c*_1_(0)*, c*_2_(0), and *c*_3_(0), respectively (Fig. 7). Using `fminsearch` in MATLAB (MathWorks, 2020), we estimate **Θ**\* such that Σ*χ*^2^ is minimised. Since *M* is discrete, while (*r, R, α, β, C*_1_(0)*, C*_2_(0)*, C*_3_(0)) are continuous, we estimate **Θ**\* for each value of *M* ∈ {0, 1, 2, 3, 4, 5} and then choose the value of *M* that minimises Σ*χ*^2^. A MATLAB implementation of this least-squares procedure is discussed in the Supplementary Information.

In Fig. 7, we show the best match that the Binary Switch Model can provide to all three datasets from Neufeld et al. (2017) for each value of *M*. The optimal parameter set **Θ**\* and minimal Σ*χ*^2^ for each value of *M* are reported in Table 6. We conclude that setting a threshold of *M* = 1 provides the best match to these datasets. While larger values of *M* clearly deviate from the experimental datasets at low population densities (Fig. 7b), setting *M* = 0 or *M* = 2 also leads to a reasonable visual match for all three experimental datasets (Fig. 7). Furthermore, it is of interest to note that the optimal model parameters associated with small values of *M* correspond to non-logistic growth features, since logistic growth can only be obtained when *r* = *R* and *α* = *β* (Table 6). The match between the experimental data and the model at *M* = 1 has several consequences: (i) this exercise confirms that the data reported by Neufeld et al. (2017) does not follow standard logistic growth; (ii) the high-quality match between the Binary Switch Model and the data for *M* = 1 is consistent with population dynamics similar to a Weak Allee effect, and; (iii) interpreting this data using the Binary Switch Model indicates that the best way to explain the population dynamics with a relatively small threshold population density.

**Table 6:** Estimates of the Binary Switch Model parameters that minimise the combined least-squres error (18) between model predicitions and experimental data from Neufeld et al. (2017). The optimal parameter set with *M* = 1, highlighted in yellow, provides the smallest combined least-squres error for all values of *M*.

| $M$ | $r$ | $R$ | $\alpha$ | $\beta$ | $C_1(0)$ | $C_2(0)$ | $C_3(0)$ | $\Sigma\chi^2$ |
| --- | --- | --- | --- | --- | --- | --- | --- | --- |
| 0 | 0.0113 | 0.0262 | 0.174 | $2.82 \times 10^{-6}$ | 0.0250 | 0.0661 | 0.184 | 0.0179 |
| 1 | 0.0168 | 0.0345 | 0.0608 | 0.0692 | 0.0192 | 0.0652 | 0.188 | 0.0154 |
| 2 | 0.0180 | 0.0576 | $2.84 \times 10^{-5}$ | 0.139 | 0.0160 | 0.0619 | 0.191 | 0.0169 |
| 3 | 0.0206 | 0.0642 | $3.66 \times 10^{-9}$ | 0.0892 | 0.0126 | 0.0534 | 0.193 | 0.0268 |
| 4 | 0.0218 | 0.134 | $3.43 \times 10^{-9}$ | 0.0623 | 0.0112 | 0.0489 | 0.191 | 0.0366 |
| 5 | 0.0237 | 0.0110 | $3.73 \times 10^{-10}$ | $2.34 \times 10^{-4}$ | 0.00933 | 0.0420 | 0.183 | 0.0571 |

## 4 Conclusions

In this work, we examine the link between threshold effects in population growth mechanisms and Allee effects. An abrupt change in growth mechanisms, which we refer to as a *binary switch*, is thought to be a common feature of biological population dynamics. Despite the ubiquitous nature of local binary switches in population dynamics, an explicit connection to Allee effects has not been considered. To explore this connection in greater detail, we examine a population density growth model, in which the proliferation and death rates vary with the local density of the population. By incorporating a local binary switch in these proliferation and death rates, we greatly reduce the size of the parameter space while explicitly incorporating a biologically realistic threshold effect in the proliferation and death rates.

To provide insight into the qualitative features of population dynamics arising in the Binary Switch Model, we examine the presence and stability of the resulting equilibria. We show that when the binary switch occurs at some intermediate population density and the high-density death rate is not too large, a diverse family of Allee effects is supported by the model. Among these Allee effects are: (i) logistic growth, when no binary switch is present; (ii) the Weak Allee effect, which modifies the simpler logistic growth model without changing its equilibria or their stability; (iii) an Extinction regime, where all population densities will eventually go extinct; (iv) the Strong Allee effect, where population below a critical density will go extinct rather than grow, and; (v) the Hyper-Allee effect, which has two distinct positive stable population densities. Furthermore, we show that there are additional forms of Allee effects at the boundaries in the parameter space that separate these five main classes of Allee effects.

Along with exhibiting a wide range of Allee effects, the Binary Switch Model has a restricted parameter regime, making the interpretation of the local binary switch clearer while requiring fewer parameters to identify when calibrating to experimental data. To demonstrate these advantages, we calibrate the Binary Switch Model to experimental datasets arising in cell biology. Not only can the Binary Switch Model provide a good match to all experimental data across three different initial densities, we also find that the parameters used to match the data provide a more explicit interpretation of the underlying local growth mechanisms arising in the population. Specifically, we confirm that the experimental data is inconsistent with the standard logistic model and that the phenomena is best explained by a binary switch at low density. We conclude that the Binary Switch Model is useful to theorists and experimentalists alike in providing insight into binary switches at the individual scale that produce Allee effects at the population scale.

While one of the merits of the Binary Switch Model is to show how a single local binary switch gives rise to a variety of Allee effects, further extensions of the modelling framework can be made. For instance, additional switches can be incorporated into the modelling framework, representing populations whose proliferation and death rates change at more than one density. We anticipate that this kind of extension would lead to additional forms of Allee effects in the resulting population dynamics. Another potential modification would be to generalise the notion how we measure local density. In this work, we take the simplest possible approach use the number of nearest neighbours on a hexagonal lattice to represent the local density. Several generalisations, such as working with next nearest neighbours or working with a weighted average of nearest neighbours, could be incorporated into our modelling framework (Fadai et al., 2019; Jin et al., 2016a). Again, we expect that such extensions would lead to an even richer family of population dynamics models. We leave these extensions for future considerations.

## Acknowledgements

This work is supported by the Australian Research Council (DP170100474).

