## Supplementary Information for "Population dynamics with threshold effects give rise to a diverse family of Allee effects"

Nabil T. Fadai[†*] and Matthew J. Simpson[‡]

[‡]School of Mathematical Sciences, Queensland University of Technology, Brisbane, Queensland 4001, Australia.

April 16, 2020

#### Contents

**S1 Binary Switch Discrete Model** **2**

**S2 Calibrating the Binary Switch Model to experimental data** **3**

### S1 Binary Switch Discrete Model

In Fig. 2, solutions of the Binary Switch Model, shown in (7), are compared with averaged simulation data of the corresponding discrete model framework. This discrete model appears in Fadai et al. (2019), whereby individuals of a population are modelled in a stochastic agent-based modelling framework. While a complete derivation of the agent-based model appears in Fadai et al. (2019), we reiterate the key features of the discrete model here.

This stochastic agent-based model, which we refer to as the Binary Switch Discrete Model, describes the temporal evolution of a population of individuals that are allowed to move, proliferate, and die based on biologically-inspired rules. We perform non-dimensional simulations on a hexagonal lattice, where each lattice site has position

$$(x_i, y_j) = \begin{cases} \left(i \ , \ j\dfrac{\sqrt{3}}{2}\right), & j \text{ even}, \\ \left(\left(i + \dfrac{1}{2}\right) \ , \ j\dfrac{\sqrt{3}}{2}\right), & j \text{ odd}, \end{cases} \tag{S1}$$

with $i = 1, \ldots, I$ and $j = 1, \ldots, J$. In Fig. 2, $I = 100$ and $J = 115$.

To incorporate crowding effects in the population, potential motility and proliferation events that would result in more than one agent per site are aborted. Agents attempt to undergo nearest neighbour motility events at rate $m_n \geq 0$, proliferation at rate $p_n \geq 0$, and death events at rate $d_n \geq 0$, where $n \in \{0, 1, \ldots, 6\}$ is the number of occupied nearest neighbour sites. For simplicity, we assume that $m_n \equiv m$ and choose $m$ such that $p_n/m \ll 1$ and $d_n/m \ll 1$ for all $n$. Agents are initially seeded on the lattice with a constant probability, representing spatially uniform initial conditions, as well as reflecting boundary conditions. We simulate the number of agents as a function of time and space using a Gillespie approach (Fadai et al., 2019) and average data from the discrete model using

$$\langle C(t) \rangle = \frac{1}{IJL} \sum_{\ell=1}^{L} Q_\ell(t). \tag{S2}$$

Here, $Q_\ell(t)$ is the total number of agents on the lattice at time $t$, in the $\ell$th identically-prepared realisation of the IBM. The total number of identically-prepared realisations is $L$; we choose $L = 100$ for the results presented in Fig. 2, in which $\langle C(t) \rangle$ is shown in red dashed curves. A description of the numerical algorithm and a MATLAB implementation of this algorithm are available at `https://github.com/nfadai/Fadai_Threshold2020`, named `BinarySwitch`.

#### S2 Calibrating the Binary Switch Model to experimental data

In Section 3, we show that the Binary Switch Model can be calibrated to experimental data by minimising the combined least-squares error between the data and model predictions. A MATLAB implementation of this calibration technique is available at `https://github.com/nfadai/Fadai_Threshold2020`, named `BinarySwitchData`. The function takes a file name of a dataset in XLS format as an input: the first column of the dataset are the time points, while the second column is the rescaled population density at these time points. `BinarySwitchData` also takes the Binary Switch Model parameters $r, R, \alpha, \beta$, and $M$ as inputs, along with the model's initial condition, $C(0)$. The function then computes the solution of (7), shown in Section 2.1, from $t = 0$ to the final time in the experimental data set. The combined least-squares error between the Binary Switch Model predictions and the experimental dataset is computed using (18). The experimental datasets shown in Neufeld et al. (2017) and Section 3 can be also found in XLS format at `https://github.com/nfadai/Fadai_Threshold2020`, named `Neufeld1`, `Neufeld2`, and `Neufeld3`.
